## Supplementary Information File for "Benchmarking DNA Foundation Models for Genomic and Genetic Tasks"

### Supplementary Note 1: Sequence classification dataset naming and source

Among the 57 sequence classification datasets, we named the datasets in a manner that clearly shows their features, and we used our naming throughout our study. Specifically:

For **4mc sites detection in multiple species**, we named the six datasets as: A.Thaliana 4mC, C.Elegans 4mC, D.Melanogaster 4mC, E.Coli 4mC, G.Pickeringii 4mC, G.Subterraneus 4mC.

For **DNase-I hypersensitive sites detection**, we named this dataset as DNase_I Hypersensitive.

For **5mC and 6mA modifications detection**, we named the two datasets as: 5-methylcytosin (5mC), N6-methyladenosine (6mA)

For **Promoter identification in multiple species**, we named the eight datasets as: Promoter GM12878, Promoter HUVEC, Promoter Hela-S3, Promoter NHEK, Promoter B_amyloliquefaciens, Promoter R_capsulatus, Promoter Arabidopsis NonTATA, Promoter Arabidopsis TATA.

For **Genomic Benchmarks Dataset Collection**, we downloaded seven datasets from https://github.com/ML-Bioinfo-CEITEC/genomic_benchmarks: demo_coding_vs_intergenomic_seqs, demo_human_or_worm, human_enhancers_cohn, human_enhancers_ensembl, human_ensembl_regulatory, human_nontata_promoters, human_ocr_ensembl. We re-named the seven datasets as: Coding, Human vs Worm, Enhancers Cohn, Enhancers Ensembl, Regulatory Region Type, Promoter NonTATA 251bps, Open chromatin region, respectively.

Besides, we have renamed the rest 33 datasets adopted from DNABERT-2 and NT-v2, in our study for better overall clarity. This document provides a mapping between the original dataset names and the nomenclature in our study. It can serve as a reference for tracing back to the original dataset sources.

**Datasets adopted from DNABERT-2**

Detailed descriptions of these datasets can be found in the original DNABERT-2 publication.

| **Our Dataset Naming** | **Original Reference** |
|---|---|
| Promoter NonTATA 300 bps<br>Promoter TATA 300 bps<br>Promoter All 300 bps | Promoter detection (Human) |
| Promoter NonTATA 70 bps<br>Promoter TATA 70 bps<br>Promoter All 70 bps | Core promoter detection (Human) |
| Human TFBS 1<br>Human TFBS 2<br>Human TFBS 3<br>Human TFBS 4<br>Human TFBS 5 | Transcription factor binding site prediction (Human) |
| Mouse TFBS 1<br>Mouse TFBS 2<br>Mouse TFBS 3<br>Mouse TFBS 4<br>Mouse TFBS 5 | Transcription factor binding site prediction (Mouse) |

| | |
|---|---|
| Yeast H3<br>Yeast H3K79me3<br>Yeast H3K9ac<br>Yeast H3K14ac<br>Yeast H3K4me3<br>Yeast H3K36me3<br>Yeast H3K4me2<br>Yeast H4<br>Yeast H3K4me1<br>Yeast H4ac | Epigenetic marks prediction (Yeast) |
| Covid variants | Covid variant prediction (Virus) |
| Splice Site Type DNABERT | Splice site prediction (Human) |

**2. Datasets adopted from NT-v2**

Detailed descriptions can be found in the corresponding sections of the original NT-v2 publication.

| **Our Dataset Naming** | **Original Reference** |
|---|---|
| Enhancer<br>Enhancer strength | Enhancer sequence prediction, Section A 4.3 |
| Splice Site Type NT<br>Donors<br>Acceptors | Splice site prediction, Section A 4.4 |

### Supplementary Note 2: Sequence classification data notes

Among the total 57 sequence classification datasets, a few of them may **not** be fully relied on experimental evidence or gold standard sources (for example, ENCODE, MethSMRT, etc.), but contains some artificial sequences. Below is the table listing the properties of each dataset. Please check the original sources of datasets for more details.

| **Dataset Name** | **is** relied on experimental evidence or gold standard sources |
|---|---|
| 5mC | Yes |
| 6mA | Yes |
| A.thaliana 4mC | Yes |
| C.elegans 4mC | Yes |
| D.melanogaster 4mC | Yes |
| E.coli_4mC | Yes |
| G.pickeringii 4mC | Yes |
| G.subterraneus 4mC | Yes |
| DNase_I Hypersensitive | Yes |
| Human TFBS 1 | Yes |
| Human TFBS 2 | Yes |
| Human TFBS 3 | Yes |
| Human TFBS 4 | Yes |
| Human TFBS 5 | Yes |
| Promoter GM12878 | Yes |
| Promoter HUVEC | Yes |
| Promoter Hela-S3 | Yes |
| Promoter NHEK | Yes |
| Promoter Arabidopsis NonTATA | Yes |
| Promoter Arabidopsis TATA | Yes |
| Promoter B_amyloliquefaciens | Yes |
| Promoter R_capsulatus | Yes |
| Acceptors | Yes |
| Coding | Yes |
| Covid Variants | Yes |
| Donors | Yes |
| Enhancer | No |
| Enhancer Strength | No |
| Enhancer Cohn | Yes |
| Enhancer Ensembl | Yes |
| Human vs Worm | Yes |
| Mouse TFBS 1 | No |
| Mouse TFBS 2 | No |
| Mouse TFBS 3 | No |
| Mouse TFBS 4 | No |
| Mouse TFBS 5 | No |
| Open Chromatin Region | Yes |

| | |
|---|---|
| Promoter All 300bps | Yes |
| Promoter All 70bps | Yes |
| Promoter NonTATA 251bps | Yes |
| Promoter NonTATA 300bps | Yes |
| Promoter NonTATA 70bps | Yes |
| Promoter TATA 300bps | Yes |
| Promoter TATA 70bps | Yes |
| Regulatory Region Type | Yes |
| Splice Site Type DNABERT | Yes |
| Splice Site Type NT | Yes |
| All Yeast Datasets | Unknown |

**Supplementary Table 1: Optimal Pooling Methods Across DNA Language Models, with one-sided Delong's Test p<0.01. Analysis based on 52 total binary sequence classification datasets.**

| ***Model*** | ***Mean Pooling*** | ***Max Pooling*** | ***Summary Token*** | ***None*** |
|---|---|---|---|---|
| *DNABERT-2* | 43 (75.0%) | 0 (0.0%) | 1 (1.9%) | 8 (15.4%) |
| *NT-v2* | 42 (80.8%) | 0 (0.0%) | 2 (3.8%) | 8 (15.4%) |
| *HyenaDNA* | 35 (67.3%) | 3 (5.8%) | 0 (0.0%) | 14 (26.9%) |
| *Caduceus-Ph* | 37 (71.2%) | 2 (3.8%) | 2 (3.8%) | 11 (21.2%) |
| *GROVER* | 41 (78.8%) | 0 (0.0%) | 0 (0.0%) | 11 (21.2%) |

**Supplementary Table 2: Baseline CNN win over DNA Foundation Models, with one-sided Delong's Test p<0.01.**

| *Dataset* | *Baseline Performs Better Than* |
|---|---|
| *5mC* | Caduceus-Ph, DNABERT-2, GROVER, HyenaDNA, NT-v2 |
| *6mA* | Caduceus-Ph, DNABERT-2, GROVER, HyenaDNA, NT-v2 |
| *A.thaliana 4mC* | Caduceus-Ph, DNABERT-2, GROVER, HyenaDNA, NT-v2 |
| *C.elegans 4mC* | Caduceus-Ph, DNABERT-2, GROVER, HyenaDNA, NT-v2 |
| *D.melanogaster 4mC* | Caduceus-Ph, DNABERT-2, GROVER, HyenaDNA, NT-v2 |
| *DNase_I Hypersensitive* | None |
| *E.coli_4mC* | Caduceus-Ph, DNABERT-2, GROVER, HyenaDNA, NT-v2 |
| *G.pickeringii 4mC* | Caduceus-Ph, DNABERT-2, GROVER, HyenaDNA, NT-v2 |
| *G.subterraneus 4mC* | Caduceus-Ph, DNABERT-2, GROVER, HyenaDNA, NT-v2 |
| *Human TFBS 1* | None |
| *Human TFBS 2* | None |
| *Human TFBS 3* | Caduceus-Ph, DNABERT-2, GROVER, HyenaDNA, NT-v2 |
| *Human TFBS 4* | DNABERT-2, HyenaDNA, NT-v2 |
| *Human TFBS 5* | None |
| *Promoter Arabidopsis NonTATA* | Caduceus-Ph, DNABERT-2, GROVER, HyenaDNA, NT-v2 |
| *Promoter Arabidopsis TATA* | Caduceus-Ph, DNABERT-2, GROVER, NT-v2 |
| *Promoter B_amyloliquefaciens* | DNABERT-2, GROVER, HyenaDNA, NT-v2 |
| *Promoter GM12878* | None |
| *Promoter HUVEC* | None |
| *Promoter Hela-S3* | None |
| *Promoter NHEK* | HyenaDNA, NT-v2 |
| *Promoter R_capsulatus* | Caduceus-Ph, DNABERT-2, NT-v2 |
| *Yeast H3* | None |
| *Yeast H3K14ac* | None |
| *Yeast H3K36me3* | None |
| *Yeast H3K4me1* | None |
| *Yeast H3K4me2* | None |
| *Yeast H3K4me3* | None |
| *Yeast H3K79me3* | HyenaDNA |
| *Yeast H3K9ac* | None |
| *Yeast H4* | HyenaDNA |
| *Yeast H4ac* | None |
| *Acceptors* | Caduceus-Ph, GROVER, HyenaDNA, NT-v2 |
| *Coding* | DNABERT-2, HyenaDNA, NT-v2 |
| *Covid Variants* | None |
| *Donors* | Caduceus-Ph, DNABERT-2, GROVER, HyenaDNA, NT-v2 |

| | |
|---|---|
| *Enhancer* | None |
| *Enhancer Strength* | None |
| *Enhancer Cohn* | HyenaDNA |
| *Enhancer Ensembl* | None |
| *Human vs Worm* | DNABERT-2, GROVER, HyenaDNA, NT-v2 |
| *Mouse TFBS 1* | Caduceus-Ph, DNABERT-2, GROVER, HyenaDNA, NT-v2 |
| *Mouse TFBS 2* | Caduceus-Ph, DNABERT-2, GROVER, HyenaDNA, NT-v2 |
| *Mouse TFBS 3* | HyenaDNA, NT-v2 |
| *Mouse TFBS 4* | Caduceus-Ph, DNABERT-2, GROVER, HyenaDNA, NT-v2 |
| *Mouse TFBS 5* | Caduceus-Ph, DNABERT-2, GROVER, HyenaDNA, NT-v2 |
| *Open Chromatin Region* | DNABERT-2, GROVER, HyenaDNA, NT-v2 |
| *Promoter All 300bps* | DNABERT-2, GROVER, HyenaDNA, NT-v2 |
| *Promoter All 70bps* | Caduceus-Ph, DNABERT-2, GROVER, HyenaDNA, NT-v2 |
| *Promoter NonTATA 251bps* | DNABERT-2, HyenaDNA, NT-v2 |
| *Promoter NonTATA 300bps* | Caduceus-Ph, DNABERT-2, GROVER, HyenaDNA, NT-v2 |
| *Promoter NonTATA 70bps* | DNABERT-2, GROVER, HyenaDNA, NT-v2 |
| *Promoter TATA 300bps* | Caduceus-Ph, DNABERT-2, GROVER, HyenaDNA, NT-v2 |
| *Promoter TATA 70bps* | Caduceus-Ph, DNABERT-2, GROVER, HyenaDNA, NT-v2 |
| *Regulatory Region Type* | None |
| *Splice Site Type DNABERT* | None |
| *Splice Site Type NT* | None |

**Supplementary Table 3: DNA Foundation Models win over baseline CNN, with one-sided Delong's Test p<0.01.**

| *Dataset* | *Models that Outperform Baseline* |
|---|---|
| *5mC* | None |
| *6mA* | None |
| *A.thaliana 4mC* | None |
| *C.elegans 4mC* | None |
| *D.melanogaster 4mC* | None |
| *DNase_I Hypersensitive* | Caduceus-Ph, DNABERT-2 |
| *E.coli_4mC* | None |
| *G.pickeringii 4mC* | None |
| *G.subterraneus 4mC* | None |
| *Human TFBS 1* | Caduceus-Ph, GROVER |
| *Human TFBS 2* | Caduceus-Ph, GROVER |
| *Human TFBS 3* | None |
| *Human TFBS 4* | Caduceus-Ph, GROVER |
| *Human TFBS 5* | Caduceus-Ph, GROVER |
| *Promoter Arabidopsis NonTATA* | None |
| *Promoter Arabidopsis TATA* | None |
| *Promoter B_amyloliquefaciens* | None |
| *Promoter GM12878* | Caduceus-Ph, DNABERT-2, GROVER, NT-v2 |
| *Promoter HUVEC* | Caduceus-Ph, DNABERT-2, GROVER, NT-v2 |
| *Promoter Hela-S3* | Caduceus-Ph, DNABERT-2, GROVER, NT-v2 |
| *Promoter NHEK* | None |
| *Promoter R_capsulatus* | None |
| *Yeast H3* | Caduceus-Ph, DNABERT-2, GROVER, HyenaDNA, NT-v2 |
| *Yeast H3K14ac* | DNABERT-2, NT-v2 |
| *Yeast H3K36me3* | Caduceus-Ph, DNABERT-2, GROVER, NT-v2 |
| *Yeast H3K4me1* | Caduceus-Ph, DNABERT-2, GROVER, HyenaDNA, NT-v2 |
| *Yeast H3K4me2* | Caduceus-Ph, DNABERT-2, GROVER, HyenaDNA, NT-v2 |
| *Yeast H3K4me3* | Caduceus-Ph, DNABERT-2, GROVER, NT-v2 |
| *Yeast H3K79me3* | Caduceus-Ph, DNABERT-2 |
| *Yeast H3K9ac* | Caduceus-Ph, DNABERT-2, GROVER |
| *Yeast H4* | Caduceus-Ph, DNABERT-2 |
| *Yeast H4ac* | Caduceus-Ph, DNABERT-2, NT-v2 |
| *Acceptors* | None |
| *Coding* | Caduceus-Ph, GROVER |
| *Covid Variants* | None |
| *Donors* | None |

| | |
|---|---|
| *Enhancer* | DNABERT-2, NT-v2 |
| *Enhancer Strength* | Caduceus-Ph, DNABERT-2, GROVER, NT-v2 |
| *Enhancer Cohn* | None |
| *Enhancer Ensembl* | None |
| *Human vs Worm* | Caduceus-Ph |
| *Mouse TFBS 1* | None |
| *Mouse TFBS 2* | None |
| *Mouse TFBS 3* | None |
| *Mouse TFBS 4* | None |
| *Mouse TFBS 5* | None |
| *Open Chromatin Region* | None |
| *Promoter All 300bps* | None |
| *Promoter All 70bps* | None |
| *Promoter NonTATA 251bps* | None |
| *Promoter NonTATA 300bps* | None |
| *Promoter NonTATA 70bps* | None |
| *Promoter TATA 300bps* | None |
| *Promoter TATA 70bps* | None |
| *Regulatory Region Type* | None |
| *Splice Site Type DNABERT* | None |
| *Splice Site Type NT* | None |

**Supplementary Table 4: The accuracy for all multi-class classification datasets in this study.** ***Bolded: row maximum with 0.01 tolerance.***

| Data | DNABERT-2 | NT-v2 | HyenaDNA | Caduceus-Ph | GROVER |
|---|---|---|---|---|---|
| Enhancer Strength | **0.715** | 0.653 | 0.69 | **0.713** | 0.703 |
| Splice Site Type, NT | 0.496 | 0.518 | **0.563** | 0.516 | 0.515 |
| Splice Site Type, DNABERT-2 | 0.605 | 0.601 | 0.618 | **0.628** | 0.611 |
| Covid Variants | **0.664** | 0.494 | 0.629 | 0.609 | **0.659** |
| Regulatory Region Type | 0.676 | 0.644 | **0.83** | 0.66 | 0.615 |

**Supplementary Table 5: Random Forest and XGBoost regression performance for gene expression prediction, averaged across all genes. P-value is comparison between random forest and XGBoost results, using two-sided Wilcoxon signed-rank test.**

| *Model* | *Input Length* | *Metric* | *Random Forest Mean* | *XGBoost Mean* | *P-value* |
|---|---|---|---|---|---|
| *DNABERT-2* | 6K bp | Correlation | **0.1208** | 0.0959 | $< 10^{-16}$ |
| | | MSE | **0.2362** | 0.2676 | $< 10^{-16}$ |
| *NT-v2* | 6K bp | Correlation | **0.1223** | 0.1038 | $< 10^{-16}$ |
| | | MSE | **0.2355** | 0.2579 | $< 10^{-16}$ |
| *HyenaDNA* | 6K bp | Correlation | **0.1224** | 0.1027 | $< 10^{-16}$ |
| | | MSE | **0.2349** | 0.2579 | $< 10^{-16}$ |
| *Caduceus-Ph* | 6K bp | Correlation | **0.1229** | 0.1024 | $< 10^{-16}$ |
| | | MSE | **0.2340** | 0.2580 | $< 10^{-16}$ |
| *GROVER* | 2K bp | Correlation | **0.1136** | 0.1035 | $< 10^{-16}$ |
| | | MSE | **0.2334** | 0.2415 | $< 10^{-16}$ |
| *Caduceus-Ph Long* | ~131K bp | Correlation | **0.1267** | 0.0947 | $< 10^{-16}$ |
| | | MSE | **0.2266** | 0.2760 | $< 10^{-16}$ |
| *HyenaDNA Long* | ~196K bp | Correlation | **0.1369** | 0.1014 | $< 10^{-16}$ |
| | | MSE | **0.2257** | 0.2700 | $< 10^{-16}$ |
| *Enformer* | ~196K bp | Correlation | **0.1294** | 0.0901 | $< 10^{-16}$ |
| | | MSE | **0.2269** | 0.2793 | $< 10^{-16}$ |

**Supplementary Table 6: Comparison of short versus long sequence models on the same set of genes. P-value is calculated using two-sided Wilcoxon signed-rank test.**

| | *Correlation (short sequence)* | *Correlation (long sequence)* | *p-value* |
|---|---|---|---|
| *HyenaDNA* | 0.122 | 0.137 | 0.0005 |
| *Caduceus-Ph* | 0.124 | 0.127 | 0.975 |

**Supplementary Table 7: Top genes ranked by random forest correlation for each model. Gene ids are in their original Ensembl ids as in GTEx dataset.**

**Short Sequence Models (2K-6K bp)**

**DNABERT-2**
ENSG00000226752.9 | 0.890423
ENSG00000214078.12 | 0.883506
ENSG00000141698.16 | 0.871117
ENSG00000013573.16 | 0.862765
ENSG00000203875.10 | 0.861914
ENSG00000280670.2 | 0.856975
ENSG00000124587.13 | 0.850042
ENSG00000006282.20 | 0.847019
ENSG00000142794.18 | 0.843301
ENSG00000172322.13 | 0.839375

**NT-v2**
ENSG00000226752.9 | 0.905441
ENSG00000258289.8 | 0.867820
ENSG00000203875.10 | 0.861324
ENSG00000124587.13 | 0.860377
ENSG00000172322.13 | 0.859079
ENSG00000013573.16 | 0.856007
ENSG00000141698.16 | 0.854827
ENSG00000280670.2 | 0.854045
ENSG00000214078.12 | 0.850859
ENSG00000229391.7 | 0.840553

**HyenaDNA**
ENSG00000226752.9 | 0.897305
ENSG00000141698.16 | 0.865093
ENSG00000214078.12 | 0.865039

ENSG00000172322.13 | 0.862775
ENSG00000258289.8 | 0.862665
ENSG00000124587.13 | 0.859704
ENSG00000280670.2 | 0.857068
ENSG00000229391.7 | 0.846921
ENSG00000006282.20 | 0.843141
ENSG00000203875.10 | 0.842432

**Caduceus-Ph**
ENSG00000226752.9 | 0.903105
ENSG00000214078.12 | 0.878051
ENSG00000141698.16 | 0.861431
ENSG00000013573.16 | 0.856631
ENSG00000258289.8 | 0.855538
ENSG00000124587.13 | 0.854781
ENSG00000172322.13 | 0.854132
ENSG00000270231.3 | 0.848280
ENSG00000280670.2 | 0.847456
ENSG00000244879.6 | 0.846667

**GROVER**
ENSG00000226752.9 | 0.892390
ENSG00000214078.12 | 0.880882
ENSG00000280670.2 | 0.863243
ENSG00000141698.16 | 0.857290
ENSG00000013573.16 | 0.856095
ENSG00000203875.10 | 0.855136
ENSG00000124587.13 | 0.855020
ENSG00000244879.6 | 0.852040
ENSG00000006282.20 | 0.850824
ENSG00000172322.13 | 0.850804

**Long Sequence Models (131K-196K bp)**

**Caduceus-Ph**
ENSG00000013573.16 | 0.830564
ENSG00000198502.5 | 0.823960
ENSG00000177697.17 | 0.777560
ENSG00000229043.2 | 0.731867
ENSG00000281741.2 | 0.728627
ENSG00000240356.6 | 0.713998
ENSG00000228716.6 | 0.694468
ENSG00000137411.16 | 0.675506
ENSG00000277402.1 | 0.644414
ENSG00000227242.4 | 0.636234

**HyenaDNA-450K**
ENSG00000013573.16 | 0.836432
ENSG00000198502.5 | 0.811447

ENSG00000177697.17 | 0.798420
ENSG00000281741.2 | 0.751786
ENSG00000229043.2 | 0.735105
ENSG00000228716.6 | 0.702603
ENSG00000240356.6 | 0.687806
ENSG00000277402.1 | 0.669714
ENSG00000253818.1 | 0.658147
ENSG00000227242.4 | 0.653490

**Enformer**
ENSG00000013573.16 | 0.841858
ENSG00000198502.5 | 0.817051
ENSG00000177697.17 | 0.813859
ENSG00000281741.2 | 0.778194
ENSG00000229043.2 | 0.751036
ENSG00000277402.1 | 0.734096
ENSG00000176393.10 | 0.718140
ENSG00000228716.6 | 0.706985
ENSG00000240356.6 | 0.695896
ENSG00000137411.16 | 0.689604

**Supplementary Table 8: The test AUC scores for each model, each test set (independent group of chromosomes) in the variant quantification effect benchmark. From up to bottom: patho, eQTL, sQTL, paQTL, ipaQTL. There are notable differences between different test sets for paQTL and ipaQTL.**

**Pathogenic variants vs common:**

| Model name | Test Group 1 AUC | Test Group 2 AUC | Test Group 3 AUC |
|---|---|---|---|
| *Caduceus-Ph* | 0.704 | 0.6877 | 0.696 |
| *Caduceus-Ph, long sequence* | 0.6296 | 0.6178 | 0.6254 |
| *DNABERT-2* | 0.5471 | 0.5322 | 0.5346 |
| *Enformer, hidden states* | 0.7145 | 0.6575 | 0.6921 |
| *Enformer, output tracks* | 0.6807 | 0.6482 | 0.6697 |
| *GROVER* | 0.6055 | 0.5942 | 0.6088 |
| *HyenaDNA-160K* | 0.624 | 0.6158 | 0.596 |
| *HyenaDNA-450K, long sequence* | 0.6357 | 0.622 | 0.6206 |
| *NT-v2* | 0.7312 | 0.7318 | 0.7326 |
| *Sei, hidden states* | 0.686 | 0.6472 | 0.6461 |
| *Sei, output tracks* | 0.6755 | 0.6495 | 0.6669 |

**eQTL:**

| Model name | Test Group 1 AUC | Test Group 2 AUC | Test Group 3 AUC |
|---|---|---|---|
| *AlphaGenome, output tracks* | 0.7925 | 0.8032 | 0.8129 |
| *Caduceus-Ph* | 0.6361 | 0.6705 | 0.641 |
| *Caduceus-Ph, long sequence* | 0.621 | 0.6441 | 0.6145 |
| *DNABERT-2* | 0.5636 | 0.5745 | 0.5723 |
| *Enformer, hidden states* | 0.7747 | 0.7706 | 0.778 |
| *Enformer, output tracks* | 0.768 | 0.7676 | 0.7742 |
| *GROVER* | 0.595 | 0.6055 | 0.5683 |
| *HyenaDNA-160K* | 0.5892 | 0.6234 | 0.6224 |
| *HyenaDNA-450K, long sequence* | 0.5914 | 0.6176 | 0.5992 |
| *NT-v2* | 0.6006 | 0.6068 | 0.62 |
| *Sei, hidden states* | 0.7434 | 0.7601 | 0.7647 |
| *Sei, output tracks* | 0.74 | 0.7604 | 0.7486 |

**sQTL:**

| Model name | Test Group 1 AUC | Test Group 2 AUC | Test Group 3 AUC |
|---|---|---|---|
| *AlphaGenome, output tracks* | 0.713 | 0.7266 | 0.7047 |
| *Caduceus-Ph* | 0.5619 | 0.541 | 0.597 |
| *Caduceus-Ph, long sequence* | 0.5755 | 0.5583 | 0.5772 |
| *DNABERT-2* | 0.5408 | 0.5984 | 0.5993 |
| *Enformer, hidden states* | 0.679 | 0.6621 | 0.6573 |

| | | | |
|---|---|---|---|
| *Enformer, output tracks* | 0.6207 | 0.6122 | 0.6192 |
| *GROVER* | 0.4652 | 0.486 | 0.4713 |
| *HyenaDNA-160K* | 0.5484 | 0.5293 | 0.5816 |
| *HyenaDNA-450K, long sequence* | 0.5306 | 0.5314 | 0.5166 |
| *NT-v2* | 0.5193 | 0.5062 | 0.4887 |
| *Sei, hidden states* | 0.6594 | 0.6168 | 0.6839 |
| *Sei, output tracks* | 0.6355 | 0.5794 | 0.668 |

**paQTL:**

| **Model name** | **Test Group 1 AUC** | **Test Group 2 AUC** | **Test Group 3 AUC** |
|---|---|---|---|
| *AlphaGenome, output tracks* | 0.6781 | 0.7354 | 0.8495 |
| *Caduceus-Ph* | 0.5059 | 0.505 | 0.5136 |
| *Caduceus-Ph, long sequence* | 0.4637 | 0.4667 | 0.4645 |
| *DNABERT-2* | 0.5118 | 0.5392 | 0.4689 |
| *Enformer, hidden states* | 0.6486 | 0.6437 | 0.7288 |
| *Enformer, output tracks* | 0.642 | 0.6275 | 0.7284 |
| *GROVER* | 0.4107 | 0.4363 | 0.5011 |
| *HyenaDNA-160K* | 0.5176 | 0.425 | 0.467 |
| *HyenaDNA-450K, long sequence* | 0.6474 | 0.5133 | 0.4954 |
| *NT-v2* | 0.5482 | 0.5821 | 0.4451 |
| *Sei, hidden states* | 0.5928 | 0.6262 | 0.6375 |
| *Sei, output tracks* | 0.621 | 0.6933 | 0.6515 |

**ipaQTL:**

| **Model name** | **Test Group 1 AUC** | **Test Group 2 AUC** | **Test Group 3 AUC** |
|---|---|---|---|
| *AlphaGenome, output tracks* | 0.8628 | 0.8525 | 0.8779 |
| *Caduceus-Ph* | 0.5712 | 0.5931 | 0.5392 |
| *Caduceus-Ph, long sequence* | 0.5278 | 0.5563 | 0.4767 |
| *DNABERT-2* | 0.4002 | 0.5619 | 0.446 |
| *Enformer, hidden states* | 0.6554 | 0.6763 | 0.7442 |
| *Enformer, output tracks* | 0.6458 | 0.6669 | 0.6634 |
| *GROVER* | 0.4549 | 0.4812 | 0.4915 |
| *HyenaDNA-160K* | 0.4332 | 0.4381 | 0.4727 |
| *HyenaDNA-450K, long sequence* | 0.4557 | 0.5775 | 0.4948 |
| *NT-v2* | 0.5686 | 0.6331 | 0.604 |
| *Sei, hidden states* | 0.6502 | 0.5813 | 0.5898 |
| *Sei, output tracks* | 0.5469 | 0.7188 | 0.5523 |

**Supplementary Table 9: HyenaDNA pretrained on DNABERT-2 multispecies dataset, compared with the HyenaDNA-1K checkpoint. Bolded: one-sided DeLong's test p-value < 0.01.**

| *Dataset* | *HyenaDNA Re-Pretrained* | *HyenaDNA-1K* |
|---|---|---|
| *5mC* | **0.7486** | 0.7065 |
| *6mA* | **0.762** | 0.7432 |
| *A.thaliana 4mC* | 0.6046 | 0.6023 |
| *C.elegans 4mC* | **0.6** | 0.5952 |
| *D.melanogaster 4mC* | 0.6222 | 0.6235 |
| *DNase_I Hypersensitive* | 0.8378 | 0.8455 |
| *E.coli_4mC* | 0.596 | 0.617 |
| *G.pickeringii 4mC* | 0.6267 | 0.622 |
| *G.subterraneus 4mC* | 0.6036 | 0.6044 |
| *Human TFBS 1* | **0.8496** | 0.8436 |
| *Human TFBS 2* | **0.8404** | 0.8308 |
| *Human TFBS 3* | 0.804 | 0.8016 |
| *Human TFBS 4* | **0.7469** | 0.7368 |
| *Human TFBS 5* | 0.9194 | 0.9223 |
| *Promoter Arabidopsis NonTATA* | 0.9066 | 0.9014 |
| *Promoter Arabidopsis TATA* | 0.7247 | 0.7245 |
| *Promoter B_amyloliquefaciens* | 0.748 | **0.7549** |
| *Promoter GM12878* | 0.7048 | 0.7058 |
| *Promoter HUVEC* | 0.6852 | 0.6912 |
| *Promoter Hela-S3* | 0.6604 | 0.6552 |
| *Promoter NHEK* | 0.8271 | 0.8351 |
| *Promoter R_capsulatus* | 0.7673 | 0.7677 |
| *Yeast H3* | 0.9113 | 0.9096 |
| *Yeast H3K14ac* | 0.7026 | 0.7083 |
| *Yeast H3K36me3* | 0.7967 | 0.7963 |
| *Yeast H3K4me1* | **0.9499** | 0.9401 |
| *Yeast H3K4me2* | 0.6428 | 0.6437 |
| *Yeast H3K4me3* | 0.8199 | 0.8108 |
| *Yeast H3K79me3* | 0.8505 | 0.8531 |
| *Yeast H3K9ac* | 0.7942 | 0.7956 |
| *Yeast H4* | 0.9398 | **0.9421** |
| *Yeast H4ac* | 0.71 | 0.715 |
| *Acceptors* | **0.9836** | 0.9682 |
| *Coding* | 0.6468 | 0.628 |
| *Covid Variants* | **0.927** | 0.9142 |
| *Donors* | **0.9216** | 0.9032 |
| *Enhancer* | 0.6563 | 0.6339 |
| *Enhancer Strength* | 0.6722 | 0.6614 |
| *Enhancer Cohn* | 0.733 | **0.7379** |
| *Enhancer Ensembl* | **0.9475** | 0.9386 |
| *Human vs Worm* | **0.8516** | 0.8326 |
| *Mouse TFBS 1* | 0.932 | 0.9297 |
| *Mouse TFBS 2* | **0.9758** | 0.9688 |

| | | |
|---|---|---|
| *Mouse TFBS 3* | **0.8748** | 0.8519 |
| *Mouse TFBS 4* | 0.7895 | 0.7821 |
| *Mouse TFBS 5* | 0.7924 | 0.7921 |
| *Open Chromatin Region* | 0.7941 | 0.8371 |
| *Promoter All 300bps* | 0.6131 | 0.6112 |
| *Promoter All 70bps* | 0.5407 | 0.5412 |

**Supplementary Table 10: Details of each sequence classification benchmark dataset.**

Table 10.1: Sequence length statistics

| Dataset Name | Median Length | Min Length | Max Length |
|---|---|---|---|
| *5mC* | 41 | 41 | 41 |
| *6mA* | 41 | 41 | 41 |
| *A.thaliana 4mC* | 41 | 41 | 41 |
| *C.elegans 4mC* | 41 | 41 | 41 |
| *D.melanogaster 4mC* | 41 | 41 | 41 |
| *DNase_I Hypersensitive* | 243 | 220 | 275 |
| *E.coli_4mC* | 41 | 41 | 41 |
| *G.pickeringii 4mC* | 41 | 41 | 41 |
| *G.subterraneus 4mC* | 41 | 41 | 41 |
| *Human TFBS 1* | 101 | 101 | 101 |
| *Human TFBS 2* | 101 | 101 | 101 |
| *Human TFBS 3* | 101 | 101 | 101 |
| *Human TFBS 4* | 101 | 101 | 101 |
| *Human TFBS 5* | 101 | 101 | 101 |
| *Promoter Arabidopsis NonTATA* | 251 | 251 | 251 |
| *Promoter Arabidopsis TATA* | 251 | 251 | 251 |
| *Promoter B_amyloliquefaciens* | 40 | 40 | 40 |
| *Promoter GM12878* | 1583 | 102 | 2995 |
| *Promoter HUVEC* | 1291 | 108 | 2993 |
| *Promoter Hela-S3* | 1109 | 101 | 2998 |
| *Promoter NHEK* | 400 | 200 | 1800 |
| *Promoter R_capsulatus* | 40 | 40 | 40 |
| *Yeast H3* | 500 | 500 | 500 |
| *Yeast H3K14ac* | 500 | 500 | 500 |
| *Yeast H3K36me3* | 500 | 500 | 500 |
| *Yeast H3K4me1* | 500 | 310 | 500 |
| *Yeast H3K4me2* | 500 | 500 | 500 |
| *Yeast H3K4me3* | 500 | 290 | 500 |
| *Yeast H3K79me3* | 500 | 500 | 500 |
| *Yeast H3K9ac* | 500 | 500 | 500 |
| *Yeast H4* | 500 | 500 | 500 |
| *Yeast H4ac* | 500 | 500 | 500 |
| *Acceptors* | 600 | 600 | 600 |
| *Coding* | 200 | 200 | 200 |
| *Covid Variants* | 999 | 999 | 999 |
| *Donors* | 600 | 600 | 600 |
| *Enhancer* | 200 | 200 | 200 |
| *Enhancer Strength* | 500 | 500 | 500 |
| *Enhancer Cohn* | 267 | 2 | 573 |
| *Enhancer Ensembl* | 200 | 200 | 200 |
| *Human vs Worm* | 200 | 200 | 200 |

| | | | |
|---|---|---|---|
| *Mouse TFBS 1* | 101 | 101 | 101 |
| *Mouse TFBS 2* | 101 | 101 | 101 |
| *Mouse TFBS 3* | 101 | 101 | 101 |
| *Mouse TFBS 4* | 101 | 101 | 101 |
| *Mouse TFBS 5* | 101 | 101 | 101 |
| *Open Chromatin Region* | 315 | 84 | 593 |
| *Promoter All 300bps* | 300 | 300 | 300 |
| *Promoter All 70bps* | 70 | 70 | 70 |
| *Promoter NonTATA 251bps* | 251 | 251 | 251 |
| *Promoter NonTATA 300bps* | 300 | 300 | 300 |
| *Promoter NonTATA 70bps* | 70 | 70 | 70 |
| *Promoter TATA 300bps* | 300 | 300 | 300 |
| *Promoter TATA 70bps* | 70 | 70 | 70 |
| *Regulatory Region Type* | 401 | 84 | 802 |
| *Splice Site Type DNABERT* | 400 | 400 | 400 |
| *Splice Site Type NT* | 400 | 400 | 400 |

Table 10.2: Sample size statistics

| **Dataset Name** | **Training Size** | **Testing Size** | **Label Distribution (Positive/Negative)** | **Label Distribution (Class 1/Class 2/…)** |
|---|---|---|---|---|
| *5mC* | 3281 | 1407 | 2344/2344 | |
| *6mA* | 25668 | 11002 | 18335/18335 | |
| *A.thaliana 4mC* | 156697 | 67157 | 111927/111927 | |
| *C.elegans 4mC* | 84926 | 36398 | 60662/60662 | |
| *D.melanogaster 4mC* | 126466 | 54200 | 90333/90333 | |
| *DNase_I Hypersensitive* | 711 | 306 | 280/737 | |
| *E.coli_4mC* | 8681 | 3721 | 2067/10335 | |
| *G.pickeringii 4mC* | 24053 | 10309 | 5727/28365 | |
| *G.subterraneus 4mC* | 63567 | 27243 | 15135/75675 | |
| *Human TFBS 1* | 24064 | 10314 | 17189/17189 | |
| *Human TFBS 2* | 22870 | 9802 | 16336/16336 | |
| *Human TFBS 3* | 14699 | 6301 | 10500/10500 | |
| *Human TFBS 4* | 20505 | 8789 | 14647/14647 | |
| *Human TFBS 5* | 14699 | 6301 | 10500/10500 | |
| *Promoter Arabidopsis NonTATA* | 8267 | 3543 | 5905/5905 | |
| *Promoter Arabidopsis TATA* | 3063 | 1313 | 1497/2879 | |
| *Promoter B_amyloliquefaciens* | 1483 | 636 | 1064/1055 | |
| *Promoter GM12878* | 10992 | 2750 | 6871/6871 | |
| *Promoter HUVEC* | 11928 | 2982 | 7455/7455 | |
| *Promoter Hela-S3* | 11736 | 2936 | 7336/7336 | |
| *Promoter NHEK* | 8170 | 2044 | 5107/5107 | |
| *Promoter R_capsulatus* | 7406 | 3175 | 5374/5207 | |
| *Yeast H3* | 10475 | 4490 | 7667/7298 | |

| | | | | |
|---|---|---|---|---|
| *Yeast H3K14ac* | 23133 | 9915 | 18771/14277 | |
| *Yeast H3K36me3* | 24415 | 10465 | 18892/15988 | |
| *Yeast H3K4me1* | 22173 | 9504 | 17266/14411 | |
| *Yeast H3K4me2* | 21478 | 9205 | 18143/12450 | |
| *Yeast H3K4me3* | 25759 | 11040 | 19604/17195 | |
| *Yeast H3K79me3* | 20185 | 8652 | 15337/13500 | |
| *Yeast H3K9ac* | 19447 | 8335 | 15415/12367 | |
| *Yeast H4* | 10220 | 4381 | 6480/8121 | |
| *Yeast H4ac* | 23866 | 10229 | 18410/15685 | |
| *Acceptors* | 15525 | 6654 | 11179/11000 | |
| *Coding* | 75000 | 25000 | 50000/50000 | |
| *Covid Variants* | 64168 | 27501 | | 11000/11000/11000/ 11000/11000/6957/ 11000/7712/11000 |
| *Donors* | 15381 | 6592 | 10973/11000 | |
| *Enhancer* | 14968 | 400 | 7684/7684 | |
| *Enhancer Strength* | 20843 | 6948 | 13895/13895 | |
| *Enhancer Cohn* | 123872 | 30970 | 77421/77421 | |
| *Enhancer Ensembl* | 14968 | 400 | | 7684/3842/3842 |
| *Human vs Worm* | 75000 | 25000 | 50000/50000 | |
| *Mouse TFBS 1* | 5668 | 2430 | 4049/4049 | |
| *Mouse TFBS 2* | 47209 | 20233 | 33721/33721 | |
| *Mouse TFBS 3* | 2293 | 983 | 1638/1638 | |
| *Mouse TFBS 4* | 1667 | 715 | 1191/1191 | |
| *Mouse TFBS 5* | 13180 | 5650 | 9415/9415 | |
| *Open Chromatin Region* | 139804 | 34952 | 87378/87378 | |
| *Promoter All 300bps* | 41437 | 17759 | 29598/29598 | |
| *Promoter All 70bps* | 41437 | 17759 | 29598/29598 | |
| *Promoter NonTATA 251bps* | 27097 | 9034 | 19657/19657 | |
| *Promoter NonTATA 300bps* | 37146 | 15920 | 26533/26333 | |
| *Promoter NonTATA 70bps* | 37146 | 15920 | 26533/26333 | |
| *Promoter TATA 300bps* | 4290 | 1840 | 3065/3065 | |
| *Promoter TATA 70bps* | 4290 | 1840 | 3065/3065 | |
| *Regulatory Region Type* | 150000 | 57713 | | 76822/68092/62799 |
| *Splice Site Type DNABERT* | 31933 | 13687 | | 10000/10000/25620 |
| *Splice Site Type NT* | 20999 | 9001 | | 10000/10000/10000 |

### Supplementary Table 11: Hyperparameter grid for gene expression prediction benchmark

**Random Forest**

N estimators: 100, 200, 500

max depth: None, 5, 10

**XGBoost**

N estimators: 100, 200, 500

Max depth: 3, 5, 7

### Supplementary Table 12: Hyperparameter grid for sequence classification benchmark

**Random Forest**

N estimators: 200, 500, 1000

max depth: None, 20

min samples split: 2,5

max features: square root, log 2

**Naïve Bayes**

Var smoothing: 1e-10. 1e-9. 1e-8

**Elastic Net Logistic Regression**

C: 0.1, 1, 10

L1 ratio: 0.1, 0.5, 0.9

class weight: balanced, None

**Supplementary Table 13: Hyperparameter grids for the random forest in variant effect quantification benchmark**

**Models with high output dimension (Enformer using either hidden states and outputs, Sei using either hidden states and outputs, AlphaGenome):**

N estimators: 100, 200, 400

Max depth: 10, None

Max features: log2, 32,

Min samples leaf: 5, 10

**Models with low output dimension (DNABERT-2, NT-v2, GROVER, Caducues-Ph, HyenaDNA, HyenaDNA-450K)**

N estimators: 100, 200, 400

Max depth: 10, None

Max features: log2, 32, sqrt

Min samples leaf: 5, 10

**Supplementary Table 14: Details of each model. The bolded ones are the included configurations in this study.**

| Model Configuration | Output Embedding Dimension |
|---|---|
| **DNABERT-2** | 768 |
| NT-v2-50m | 512 |
| NT-v2-100m | 512 |
| NT-v2-250m | 768 |
| **NT-v2-500m** | 1024 |
| Hyena-tiny-1k | 128 |
| **Hyena-tiny-1k-d256** | 256 |
| Hyena-tiny-16k-d128 | 128 |
| Hyena-small-32k | 256 |
| **Hyena-medium-160k** | 256 |
| **Hyena-medium-450k** | 256 |
| Hyena-large-1m | 256 |
| **GROVER** | 768 |
| **Caduceus-Ph-131k-d-256** | 256 |
| Caduceus-Ph-1k-d-256 | 256 |
| Caduceus-Ph-1k-d-128 | 118 |

**Supplementary Figure 1: Classifier comparison, for all models and pooling methods.** We calculate AUC scores from all 52 binary sequence classification datasets included in this study. Boxplots show the median (center line), interquartile range (box = 25th–75th percentiles), and whiskers correspond to 1.5 × interquartile range. minima and maxima correspond to the whisker ends.

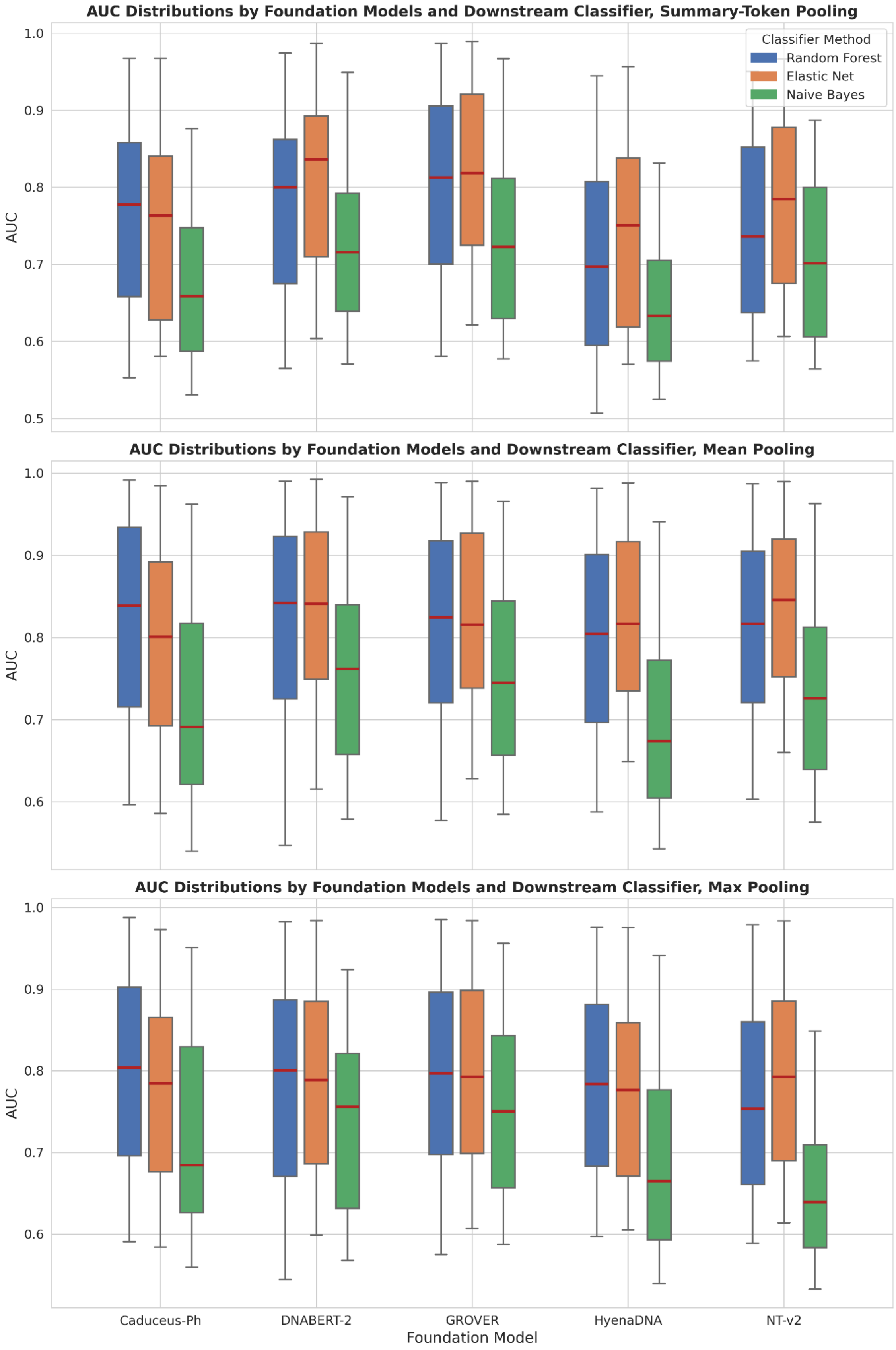

**Supplementary Figure 2: The heatmap of the difference between attention matrices for TAD-centered versus background sequences.** The horizontal axis represents key tokens, and the vertical axis represents query tokens. Each point (x,y) shows the difference in attention weight that query token y places on key token x when comparing TAD-centered versus background sequences. The central 400 tokens (approximately positions 200-600) correspond to the TAD region. If NT-v2 recognized TAD boundaries, we would expect a vertical band of positive differences in this central region.

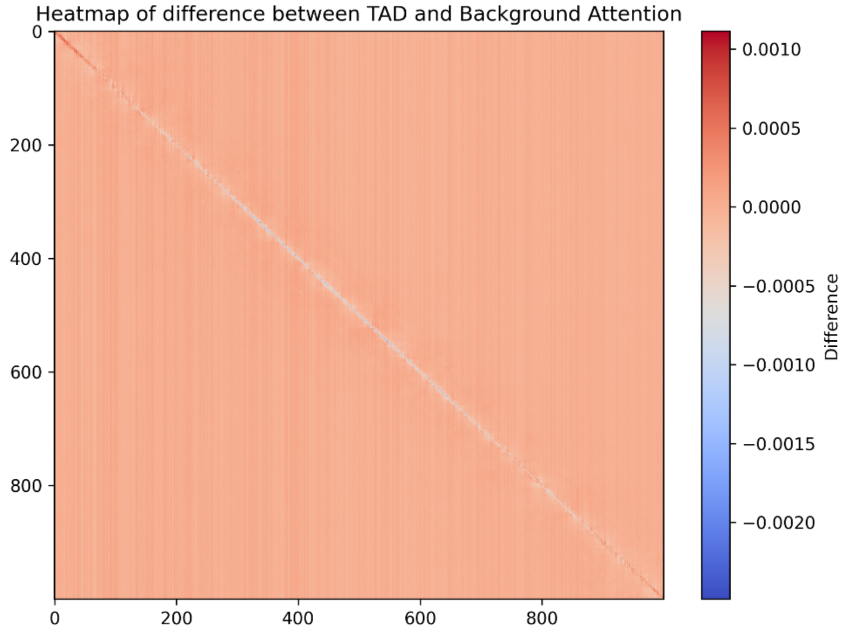
